# Terminal nucleotidyltransferase *Tent2* microRNA tailing regulates excitatory/inhibitory balance in the hippocampus

**DOI:** 10.1101/2024.08.22.609125

**Authors:** P. Wardaszka, B. Kuzniewska, N. Gumińska, A. Hojka-Osinska, M. Puchalska, J. Milek, A. Stawikowska, P. Krawczyk, F.P. Pauzin, T. Wojtowicz, K. Radwanska, C. Bramham, A. Dziembowski, M. Dziembowska

## Abstract

One of the post-transcriptional mechanisms regulating the stability of RNA molecules involves the addition of non-templated nucleotides to their 3’ ends, a process known as RNA tailing. To systematically investigate the physiological consequences of terminal nucleotidyltransferase TENT2 absence on RNA 3’ end modifications in the mouse hippocampus we developed a new *Tent2* knockout mouse. Electrophysiological measurements revealed increased excitability in *Tent2* KO hippocampal neurons, and behavioral analyses showed decreased anxiety and improved fear extinction in these mice. At the molecular level, we observed a significant contribution of TENT2 to the monoadenylation of various classes of miRNAs, but found no effect of the enzyme’s loss on the total poly(A) tail length of mRNAs, as measured by Direct Nanopore RNA sequencing. Alterations in monoadenylation of a large population of microRNAs affected the overall mRNA abundance, particularly transcripts related synaptic transmission, which were downregulated in the hippocampus of *Tent2* knockout mice. These changes explain the observed behavioral and electrophysiological alterations. Our data thus establish a link between TENT2-dependent microRNA tailing and the balance of inhibitory and excitatory neurotransmission.

## Introduction

The mRNA poly(A) tails added by canonical nuclear PAP in the nucleus are vital for regulating mRNA stability, export to the cytoplasm, translation, and decay. In addition to classical poly(A) adenylation, the 3’ ends of various classes of RNAs are modified by enzymes known as terminal nucleotidyltransferases (TENTs), which add non-templated nucleotides to their 3’ ends, a process known as RNA tailing. These post-transcriptional modifications affect RNA stability and function and were shown to be crucial in early embryonic development and gamete production. Cytoplasmic polyadenylation is best known from studies of gametogenesis (J. H. Kim & Richter, 2006, K. W. Kim et al., 2010, Sartain et al., 2011, Wang et al., 2002) where specific mRNAs are rapidly polyadenylated in response to cellular signals, allowing translation to start in a transcription-independent fashion.

In humans, there are 11 different terminal nucleotidyltransferases (TENTs), that are enzymes responsible for the posttranscriptional addition of nontemplated nucleotides to the 3′ end of RNA molecules. TENT2 (terminal nucleotidyltransferase 2, GLD2) is a noncanonical poly(A) polymerase, that adds adenine nucleotides to the 3’ end of RNAs. TENT2 was first described in non-mammalian species, including *C. elegans*, *X. laevis* and *Drosophila melanogaster*. In these organisms, TENT2 was shown to elongate poly(A) tails of a subset of stored cytoplasmic mRNAs, thus regulating thei translational activation in early development (Nakel et al., 2015, Wang et al., 2002, Nousch et al., 2017, Barnard et al., 2004, Rouhana et al., 2005). This process, known as cytoplasmic polyadenylation, is a major regulatory mechanism during oocyte maturation and egg activation (Cui et al., 2013).

In contrast, in mice, *Tent2* disruption did not affect the maturation of the oocytes or animals’ fertility (Nakanishi et al. 2007), nor the poly(A)tail elongation in oocytes using reporter RNAs. *Tent2*-deficient mice remain fertile and healthy, suggesting that other TENTs may be involved in mRNA polyadenylation in the mammalian germline. Interestingly, TENT2 is widely expressed in the mouse brain, where it can regulate posttranscriptional modifications of RNAs involved in synaptic plasticity. TENT2 was suggested to polyadenylate specific target mRNAs, such as NR2A subunit of N-methyl-D-aspartate receptor (Udagawa et al. 2012), essential for long-term synaptic plasticity. However, *Tent2* knockout mice generated with a classical ES call-based approach do not display any obvious behavioral abnormalities (Mansur et al. 2016).

In addition to its role in mRNA expression, TENT2 was shown to regulate the stability of certain mature miRNAs, and its deletion in a human fibroblast cell line significantly reduced the fraction of tailed miRNAs possessing an untemplated monoadenosine (D’Ambrogio et al. 2012). The liver-specific miRNA-122, is 3′ monoadenylated in human hepatocytes and mouse liver (Katoh et al. 2009). TENT2 knockout mice show significantly lower levels of miRNA-122, indicating that TENT2 monoadenylation enhances its stability (Katoh et al. 2009).

Knowing that *Tent2* is ubiquitously expressed in the nervous system, in this study, we generated a *Tent2* knockout mouse line using CRISPR/Cas9 to revisit the potential role of this enzyme in neuronal RNA metabolism and physiology. We observed a significant contribution of TENT2 to the monoadenylation of various classes of miRNAs, but found no effect of the enzyme’s loss on the total poly(A) tail length of mRNAs, as measured by direct Nanopore sequencing. The alterations in monoadenylation of a large population of microRNAs apparently affected the overall mRNA abundance. Specifically, we observed a group of transcripts related to neurotransmitter transport and synaptic transmission that were downregulated in the hippocampus of *Tent2* knockout mice. The electrophysiological measurements revealed increased excitability of *Tent2* KO neurons, and the analyses of mice behavior have shown decreased anxiety and improvement in fear extinction of *Tent2* KO mice. Our data establish a link between TENT2-dependent microRNA tailing and the balance of inhibitory and excitatory neurotransmission.

## Results and Discussion

To assess the role of TENT2 poly(A) polymerase in the brain, we generated a new TENT2 knockout mouse line using genome editing technology (Fig. 1 A). The homozygous *Tent2* KO mice displayed no visibly harmful phenotype, were viable and fertile, and produced similar litter sizes to wild-type mice. To elucidate the physiological consequences of the absence of TENT2 enzymatic activity on RNA 3’ end modifications in the mouse hippocampus, we conducted: i) microRNA sequencing, ii) poly(A) tail length analysis using direct nanopore sequencing, iii) total mRNA sequencing with Illumina.

**Figure 1.**
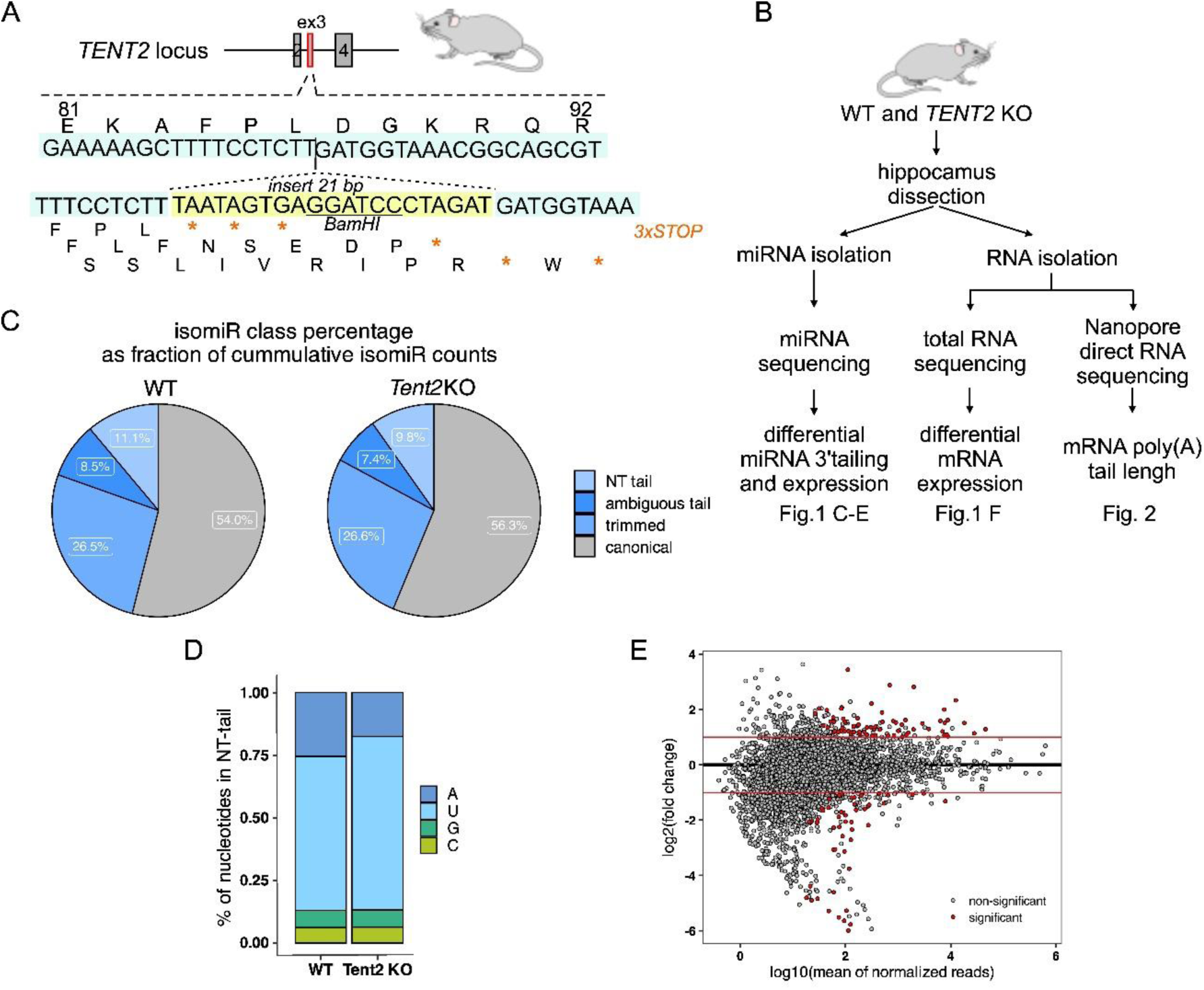
miRNA sequencing and poly(A) tail length analysis in the hippocampus of *Tent2* KO mice. **A**, Schematic illustration of *Tent2* locus fragment. Position of the 21bp insert is indicated, the sequence is marked with a yellow box. To facilitate the detection of mutations, a restriction site for BamHI was introduced. STOP codons introduced in the insert are marked with asterisks. **B**, Workflow of the experiments presented in panels C-F and on Figure 2, depicting isolation of RNA from the hippocampi and different sequencing approaches. **C-E**, Results of miRNA sequencing of *Tent2* KO and WT hippocampi (WT n=3, *Tent2* KO n=5). **C**, IsomiR class percentage analysis, showing that 11,1% of isomiRs in WT and 9,8 % in *Tent2* KO had NT tail at the 3’ end. **D,** Significant reduction of adenylation accompanied by increase in uridylation in the NT tails in *Tent2* KO hippocampi. **E,** miRNA abundance in *Tent2* KO hippocampi as compared to WT.

### TENT2-dependent adenylation of miRNA does not affect their stability in mouse hippocampus

In the previous study of Mansur et al. the authors have shown a reduction in the amount of monoadenylation of miRNAs, but not in the miRNA levels themselves in the hippocampus of another *Tent2* KO mouse model (Mansur et al. 2016). To corroborate this study in our *Tent2* KO mice and further correlate the results with mRNA polyadenylation status we performed small RNA Sequencing (sRNA-Seq). We generated the sRNA libraries using hippocampi from 5 *Tent2* KO and 3 wild-type (WT) male mice (littermates). As our focus was the 3’ non-templated additions to the miRNAs we analyzed miRNA expression levels and 3’ miRNA isoforms (isomiRs) composition using QuagmiR (Bofill-De Ros et al. 2019). It is a highly customizable tool for comprehensive analysis of heterogeneous isomiRs populations and allows for various 5’ and 3’ templated and not templated (NT), tailed as well as trimmed isomiRs detection. We consistently detected 3452 isomiRs resulting from 576 miRNAs in all 8 analyzed samples. The primary analysis showed that more than 50% of analyzed reads were assigned to the canonical form of miRNA (annotated mature miRNA sequence). Then, on average, 26.5% of reads represented trimmed miRNA isoforms, and the remaining were tailed isomiRs. According to our results, on average, only 10% of isomiRs had an NT tail at their 3’ end. As expected for the tailing enzyme, *Tent2* KO leads to reduced accumulation of isomiRs with NT-tails (Fig. 1 C and Fig. S1A). Notably, adenosine frequency is significantly reduced in the NT tails under *Tent2* KO (Fig. 1 D and Fig. S1B). The decreased adenylation was accompanied by a moderate increase in uridylation, which can explain why the drop on the overall NT tail was not as deep as one may expect.

Further, we analyzed NT tails in miRNAs by their length and composition. Intriguingly, we did not observe a significant decrease in global 1 nucleotide (nt) additions but only in the 2 and 3 nt additions (Fig. S1C). This was explained when the changes in tail composition by length were examined. *Tent2* KO led to a reduction of accumulation of any type of tail that contained adenosines. The decrease of mono-A tails was compensated by the mono-U and mono-C additions. For the 2 and 3 nt tails we observed a prominent drop in the accumulation of tails containing adenosines.

To explore the impact of reduced mono-adenylation on miRNA abundance, we compared the levels of all isomiRs in *Tent2* KO hippocampi with the WT. The analysis revealed that more than 7% of isomiRs were significantly dysregulated in *Tent2* KO, with a similar number of up- and down-regulated ones (Fig. 1 E and Supplementary Table 1 and 2). The differential analysis confirmed our observations of changes in tail composition, as the pattern of drop of many isomiRs under the *Tent2* KO is mainly due to decreased accumulation of isomiRs with tails enriched in adenosines (Fig. S1D). As the tailed isomiRs are not the main form of miRNA expressed, we examined how the change in accumulation of mono-A tail isomiRs (as an example of TENT2 action) correlates with the accumulation of canonical form. This analysis showed no correlation between the changes in 3′ mono-adenylation and the alteration in abundance of canonical miRNA (Fig. S1E). The fact that both upregulated and downregulated canonical miRNAs were observed in *Tent2* KO samples when the mono-A tailed form of miRNAs was decreased indicates that the underlying mechanisms are complex. Together, these results show that TENT2 is a primary enzyme responsible for adenylation of miRNAs in mouse hippocampus and that the TENT2 mediated adenosine addition does not globally affect the main miRNA isoforms stability.

### *Tent2* knockout mice show changes in expression of mRNAs related to neurotransmission and no differences in mRNA poly(A) tail length

In addition to miRNA analysis, the RNA samples isolated from *Tent2* KO and WT hippocampi were processed using two high-throughput platforms. The mRNA expression levels were determined by total RNA sequencing with Illumina (RNA-seq), whereas the poly(A) tail lengths were assessed at the whole transcriptome scale by Oxford Nanopore Direct RNA Sequencing (DRS) (Fig. 1 B).

Differential expression analysis of the RNA-seq data, provided in Supplementary Table 2, identified 134 differentially expressed genes: 66 downregulated and 68 upregulated Fig. 2 A). Notably, a majority of the upregulated genes (44) are annotated as putative, mostly regulatory *loci*, derived from automated computational methods. Whereas all the downregulated mRNAs correspond to well-characterized, reliable protein-coding genes. Consequently, our focus shifted toward these downregulated genes. Gene ontology analysis revealed a significant overrepresentation of mRNAs involved in the regulation of membrane potential, neurotransmitter transport, glutamatergic synaptic transmission, learning and memory, and inhibition of synaptic transmission among genes downregulated in *Tent2* KO mice (Fig. 2 B).

**Figure 2.**
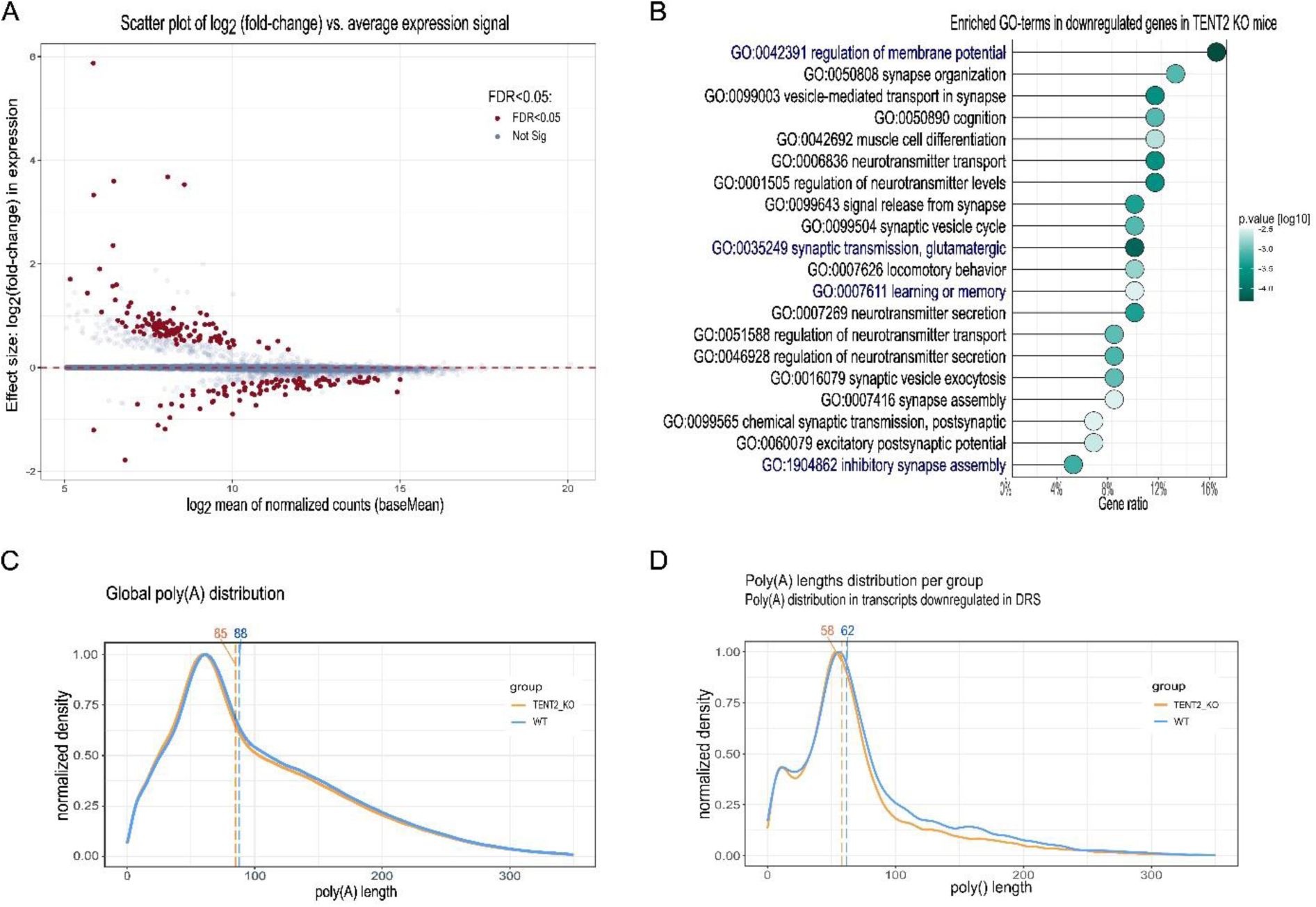
Results of RNA sequencing of mRNAs in the hippocampus of *Tent2* KO and WT mice. **A,** Differential expression of mRNAs in *Tent2* KO hippocampi vs. WT (n=3/genotype). Genes with significantly changed expression are highlighted (red). **B,** Gene ontology analysis of transcripts downregulated in *Tent2* KO mice. **C, D,** Poly(A) tail length analysis of mRNAs in the hippocampus of *Tent2* KO and WT mice (n=3 mice/genotype). No differences in global mRNA poly(A) tail length (C) and transcripts with altered expression (D) were detected in *Tent2* KO mice as compared to WT.

We then examined the poly(A) tail lengths of the DRS data. Interestingly, we found no global effect in poly(A) tail length distributions between the studied phenotypes (Fig. 2 C and S2A). Moreover, we did not observe significant changes in poly(A) tail lengths in genes with altered expression either (both down- and upregulated) (Fig. 2 D and S2B).

### Increased excitability and decreased inhibitory synaptic transmission in *Tent2* KO hippocampal neurons

Among the mRNAs differentially expressed in *Tent2* KO hippocampus, we identified an overrepresentation of transcripts involved in regulation of neurotransmitter levels and synaptic transmission. We, therefore, aimed to assess the electrophysiological properties of neurons from *Tent2* KO using patch-clamp. Recordings were performed in acute brain slices prepared from 6 WT and 4 *Tent2* KO mice. We measured passive membrane properties and membrane excitability of pyramidal neurons in the CA1 hippocampal region (Fig. 3 A). Rheobase, the minimal current magnitude that evoked an action potential, was significantly decreased in *Tent2* KOs (Fig. 3 B-C; Mann-Whitney test; * p = 0.0257). The majority of *Tent2* KO neurons generated an action potential at 25 pA of applied current, whereas most of the WT neurons started firing at 50 pA. First spike latency (Fig. S3A), general frequency of spiking (Fig. S3B) and accommodation ratio (Fig. S3C) remained unchanged but the frequency of the spikes during the first half of the stimulus pulse was decreased for *Tent2* KO neurons at final current values. We did not observe any significant differences in membrane capacitance (Fig. 3 D), membrane resistance (Fig. 3 E) or resting membrane potential (Fig. 3 F) between *Tent2* KO and WT neurons. Altogether, *Tent2* KO neurons showed increased excitability in response to current stimuli without a change in the key passive membrane properties or kinetics of individual action potentials.

**Figure 3.**
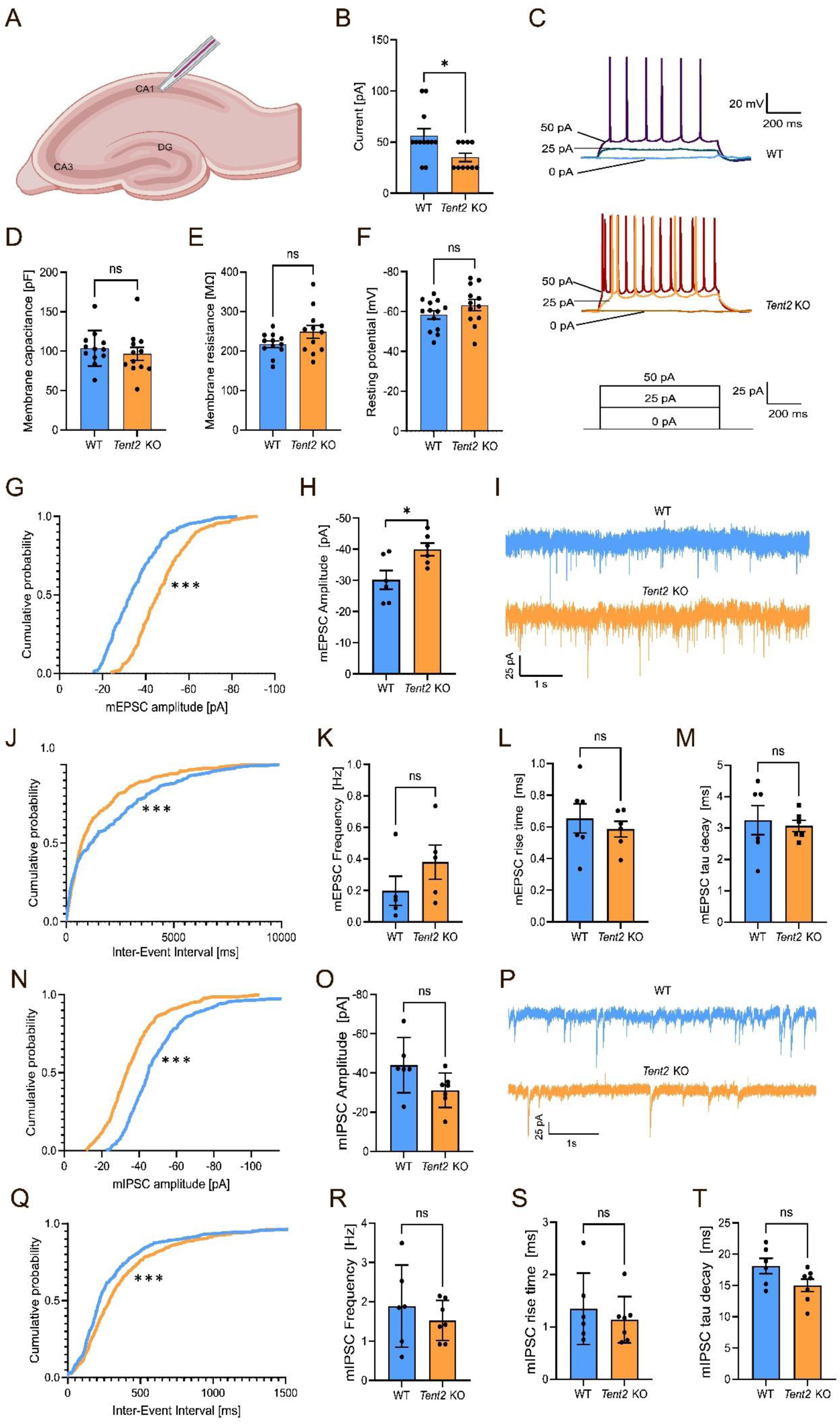
*Tent2* KO neurons exhibit increased excitability in CA1 in hippocampal slices (A-F) and enhanced excitatory synaptic transmission in cultured hippocampal neurons (G-M) **A,** Schematic of patch-clamp recordings in CA1 region of hippocampal horizontal slices. **B,** Rheobase, the lowest value of current to trigger an action potential was significantly lower for *Tent2* KO (Mann-Whitney test; * p = 0.0257). **C,** Representative voltage responses to current stimuli for WT (top panel) and *Tent2* KO neurons (middle panel). **D-F,** Statistics for selected passive membrane properties (see Methods for description). Membrane capacitance; membrane resistance or resting membrane potential were not altered in Tent2 KO neurons. Data were obtained from 12-13 neurons per group. The data are expressed as mean +/- SEM. **G-M,** AMPA/kainate receptor glutamatergic transmission was recorded in 21-22 DIV hippocampal cultures with whole-cell patch-clamp method. **G,** The cumulative probability of individual mEPSCs amplitudes was shifted toward higher values (Kolmogorov-Smirnov test; p<0.00001). **H,** Averaged amplitude of mEPSCs was significantly larger for Tent2 KO neurons (t-test; p<0.05). **I,** Representative current traces from WT and Tent2 KO neurons. **J,** The cumulative distribution function for inter-event time intervals between subsequent mEPSCs is left-shifted compared to WT neurons. (Kolmogorov-Smirnov test P<0.0001). **K,** The averaged frequency of mEPSCs is not significantly changed between genotypes. L-M, *Tent2* KO neurons do not exhibit altered mEPSCs rise time (**L**) or tau decay (**M**). Data were obtained from 6 neurons per group. The data are expressed as mean +/- SEM. **(N-T)** Reduction of inhibitory synaptic transmission in *Tent2* KO cultured hippocampal neurons. **N**, The cumulative probability of amplitudes for mIPSCs is left-shifted (Kolmogorov-Smirnov test; p<0.0001). **O**, Averaged amplitudes of mIPSCs were not significantly different in *Tent2* KO neurons (t-test; p=0.07). **P**, Representative current traces of mIPSCs recordings from WT and *Tent2* KO cultured neurons (21-22 DIV). **Q**, The cumulative probability plot of the inter-event interval for *Tent2* KO is right-shifted relative to the WT neurons (Kolmogorov-Smirnov test P<0.0001). **R**, The averaged frequency of mIPSCs was not significantly changed. **S-T**, mIPSCs rise time **(S)** and tau decay **(T)** were not altered in *Tent2* KO neurons. Data were obtained from 6-7 neurons per group. The data are expressed as mean +/- SEM. Partially created with Biorender.

Increased excitability of pyramidal cells in hippocampal CA1 may be due to enhanced excitatory or reduced inhibitory synaptic transmission. In order to verify which one is disturbed, we measured both miniature excitatory and inhibitory currents with the whole-cell patch-clamp method. To this end, we chose to study cultured neurons, which exhibit a significant amount of synaptic events compared to the acute brain slices model. First, we measured miniature excitatory postsynaptic currents (mEPSCs) in 21-22 DIV-cultured primary hippocampal neurons. In *Tent2* KO neurons, the amplitude of mEPSCs was increased, as shown in the cumulative probability of amplitudes for individually recorded miniature currents (Fig. 3 G; Kolmogorov-Smirnov test; p<0.00001) and the averaged mEPSCs amplitudes (Fig. 3 H; t-test; p<0.05). The representative traces (Fig. 3 I) suggested increased mEPSC frequency, and this was corroborated by the cumulative distribution function showing a significantly shorter mEPSC inter-event interval in *Tent2* KO neurons relative to WT (Fig. 3 J; Kolmogorov-Smirnov test P<0.0001). In addition, averaged mEPSCs frequency (Fig. 3 K) was almost two times higher in *Tent2* KO neurons, although this change was not statistically significant. The rise-time (Fig. 3 L) and tau decay (Fig. 3 M) of the currents were also not significantly different between genotypes. Thus, cultured *Tent2* KO neurons exhibited enhanced excitatory synaptic transmission. These results confirm the results obtained with acute brain slices.

Next, we recorded GABA_A_ receptor-mediated miniature inhibitory postsynaptic currents (mIPSCs) in 21-22 DIV cultured primary hippocampal neurons. The comparison of cumulative probability of mIPSCs amplitude between WT and *Tent2* KO (Fig. 3 N) neurons showed that mIPSCs amplitudes are smaller in *Tent2* KO neurons (Kolmogorov-Smirnov test; *** p<0.0001). Averaged amplitudes (Fig. 3 O) demonstrated the same trend but the difference was not statistically significant (t-test; p=0.07). The intervals between subsequent mIPSCs (Fig. 3 Q) was elongated in *Tent2* KO neurons (Kolmogorov-Smirnov test; *** P<0.0001). However, there was no visible reduction of the average mIPSC frequency (Fig. 4 R).

**Figure 4.**
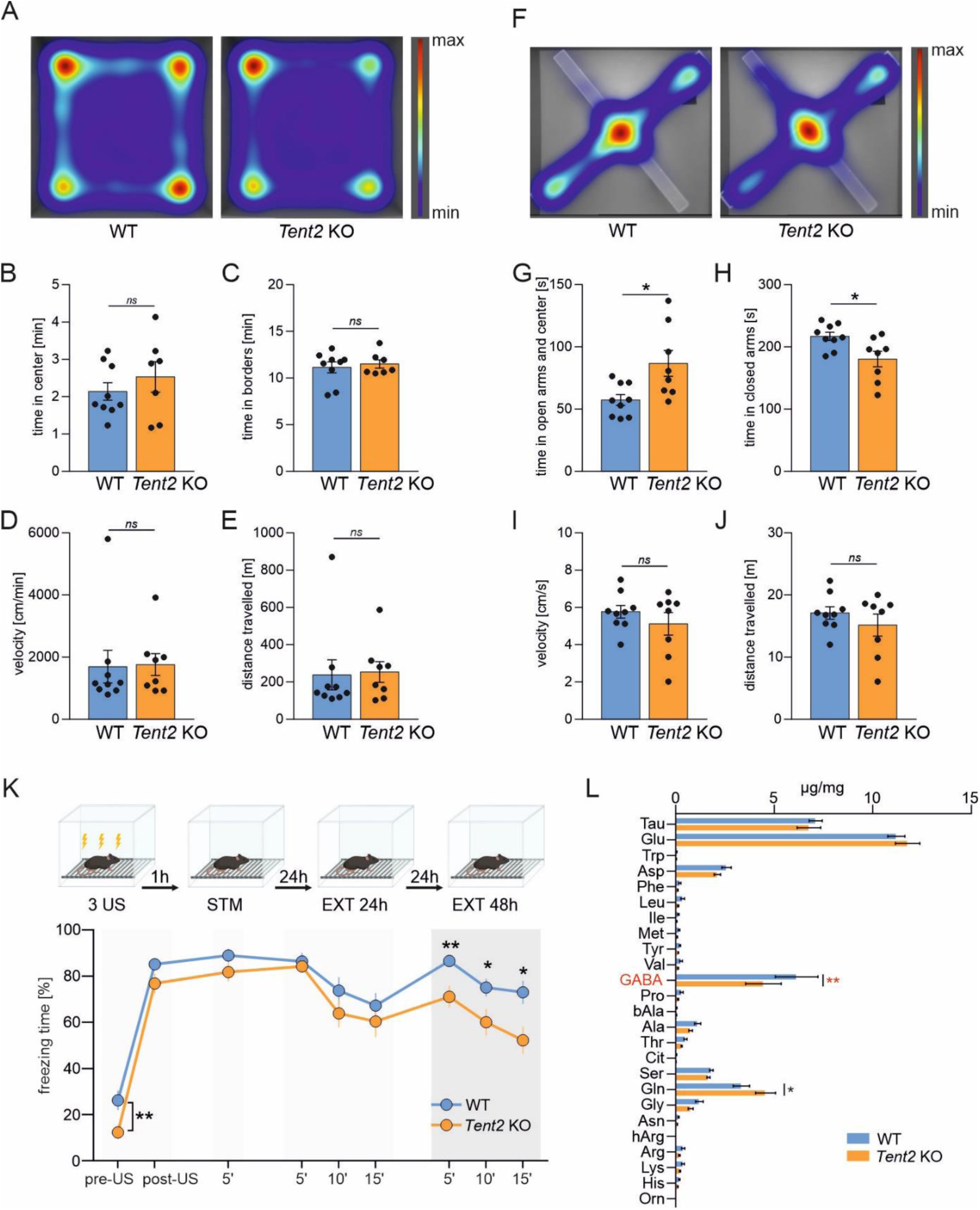
Behavioral characterization of *Tent2* KO and WT mice. Anxiety, exploratory behavior and locomotion was assessed using open field (A-E) and elevated plus maze (F-J) tasks. Learning and memory was assessed using contextual fear conditioning (K). **A**, Population averaged heatmaps showing mice activity in the open arena. No significant differences in time spent in the center (B), time spent in borders (C), velocity (D) or distance moved (E) were observed (p-value > 0.05, unpaired t-test, n=8-9). F, Population averaged heatmaps showing mice activity in the elevated plus maze. *Tent2* KO mice spent significantly more time in the open arms and in the center as compared to WTs (G, * p-value = 0.0156, unpaired t-test, n=8-9) and significantly less time in the closed arms (H, p-value = 0.017, unpaired t-test, n=8-9). Similarly as in the open field task, no significant differences in velocity (I) or distance traveled (J) were observed (p-value > 0.05, unpaired t-test, n=8-9). **K**, Experimental timeline (upper panel) and summary of data showing freezing levels during contextual fear conditioning (CFC) and test sessions. Mice were conditioned in a novel context (3US presentations, 2s, 0.7 mA). Next, the mice were re-exposed three times to the same context: 1 hour after CFC to assess short-term contextual fear memory (STM, 5 min); 24 hours after CFC to test long-term contextual fear memory and to extinguish contextual fear (EXT1, 15 min); and 48 hours after CFC to test fear extinction memory (EXT2, 15 min). *Tent2* KO mice showed increased locomotion before the training (pre-US) as compared to WT animals (**, p-value = 0.01), but did not differ in freezing behavior post-US (p-value > 0.05), 1h after training (STM) nor 24 hours after training (LTM). Interestingly, *Tent2* KO mice showed decreased freezing levels during EXT2 (** p-value = 0.009, * p-value = 0.039, * p-value = 0.012). Data is presented as mean +/- SEM; n = 18-22; Two-way ANOVA, post-hoc Sidak’s multiple comparisons test. **L**, Levels of amino acids in the hippocampi of fear-conditioned *Tent2* KO and WT mice. Directly after the last test in the contextual fear conditioning experiment shown in panel K, mice were sacrificed and the levels of amino acids was measured using mass spectrometry. Significantly decreased levels of GABA neurotransmitter were detected in the hippocampi of *Tent2* KO mice as compared to WT (** p-value = 0.0038). Additionally, increased levels of glutamine were observed in *Tent2* KO hippocampi (* p-value = 0.0267). Data is presented as mean +/- SEM; n = 8-9; Two-way ANOVA, post-hoc Sidak’s multiple comparisons test.

Altogether, electrophysiological measurements in hippocampal neurons indicated that *Tent2* KO neurons may exhibit increased excitability. In addition, when *Tent2* KO neurons are cultured, they exhibit enhanced excitatory and decreased inhibitory synaptic transmission. Therefore, we have next examined whether observed alterations in excitatory/inhibitory balance following *Tent2* KO may be associated with altered behavior in mice.

### Decreased anxiety and increased fear memory extinction in *Tent2* KO mice

We performed behavioral tests to assess anxiety, locomotor activity as well as learning and memory. Firstly, we tested the mice in the open field task (Fig. 4 A-E). We did not observe any significant differences in the locomotor activity (velocity and distance travelled, Fig. D, E) of *Tent2* KO and WT mice. Similarly, we did not detect any differences in the marble burying test (Fig. S4), that is used to study compulsive-like behaviors. In contrast, we observed increased exploration of open arms and decreased time spent in closed arms of elevated plus maze by *Tent2* KO mice (Fig. 4 F-J), indicating decreased anxiety. Similarly as in the previous tests, no significant differences in the locomotor activity were observed (Fig. 4 I-J). Our data is consistent with the results of (Mansur et al. 2016) where in another *Tent2* KO model no behavioral differences in the open field and marble burying tests were seen. One significant difference in our study was decreased anxiety of *Tent2* KO mice in elevated plus maze, which was not observed by Mansur and colleagues.

Since *Tent2* KO mice exhibited decreased anxiety, we decided to check if they will show differences in contextual fear memory acquisition and extinction (Fig. 4 K). We noticed an increase in locomotion (low freezing level) in *Tent2* KO mice before the training (p-value = 0.01). This difference disappeared after delivery of 3 electric shocks (3US). There was no difference between WT and *Tent2* KO mice in freezing frequency during short-term (STM) and long-term memory test (EXT 24h), indicating no impairments in formation and expression of short- and long-term memory. Moreover, when animals were exposed to the training context for 15 minutes without US presentation for contextual fear memory extinction, both WT and *Tent2* KO mice decreased freezing levels within the session and no difference between the genotypes were observed (p-value = 0.009, p-value = 0.039, p-value = 0.012). However, when fear extinction memory was tested on the next day, freezing levels were higher in WT mice, indicating that fear memory extinction was increased in *Tent2* KO mice.

### Decreased level of GABA in the hippocampus of *Tent2* KO

As we found changes in anxiety and fear extinction memory in *Tent2* KO mice, alterations in excitatory/inhibitory balance as well as downregulation of transcripts involved in neurotransmitter transport and synaptic transmission, we decided to measure the concentration of amino acids and neurotransmitters in the hippocampus of *Tent2* KO and WT mice. After the last fear extinction test the level of GABA, a principal inhibitory neurotransmitter in the central nervous system, was significantly reduced (p-value = 0.0038) in *Tent2* KO. Moreover, concentration of glutamine, that is a precursor for glutamate, was increased (p-value = 0.0267). Although we did not observe statistically significant differences in the glutamate levels themselves, there was a trend towards its increased levels. These data align with decreased GABA-ergic neurotransmission and increased excitability in *Tent2* KO neurons (Fig. 3).

Our data suggest that the excitatory/inhibitory (E/I) balance is disturbed in Tent2 KO mice. Based on electrophysiological measurements we observed increased intrinsic membrane excitability with no alteration in membrane potential or resistance that could explain this observation. Therefore we focused our attention on other factors that may contribute to the alternation of E/I balance and we observed a significant increase in miniature excitatory and a decrease in inhibitory postsynaptic currents. Similar results were observed in a mouse model of tuberous sclerosis (TSC1 deficiency), where hyperactivity of neuronal network was caused by a decrease in the amplitude and frequency of inhibitory miniature currents (Bateup et al. 2013).

Functioning of both principal excitatory neurons as well as inhibitory interneurons is crucial for maintaining the physiological balance of excitatory and inhibitory neurotransmission in the brain. Consequently, the dysregulation of E/I balance is a common feature of different neurological and neuropsychiatric disorders such as anxiety (Kalueff and Nutt 2007), schizophrenia (Heckers & Konradi, 2015, Nguyen et al., 2014) bipolar disorder (Brady et al. 2013) and autism spectrum disorder (ASD) (Pizzarelli and Cherubini 2011). Actually, one of the ASD proposed mechanisms is the E/I imbalance theory. According to this theory reduced inhibition in the cortex and hippocampus leads to over-stimulation of neurons and less efficient information processing (Rubenstein & Merzenich, 2003, Sohal & Rubenstein, 2019). Reduced inhibition was also reported in Mecp2 KO mice, a model of Rett syndrome and autism spectrum disorder. GABAergic neurons lacking MECP2 exhibit a decrease in GABA release and reduced level of mRNAs encoding GAD65 and 67 (Chao et al. 2010). In our *Tent2* KO mice we detected decreased amplitude and increased inter-event intervals when the miniature inhibitory postsynaptic currents were measured, which indicates reduced GABAA receptor content per synapse and a lower number of inhibitory synapses. The level of GABA, the main inhibitory neurotransmitter in the CNS was significantly reduced in *Tent2* KO mice after fear conditioning. Our data support the role of hippocampal GABAergic circuits in the regulation of fear acquisition (Oh and Han 2020). Abnormal hippocampal disinhibition may lead to attentional and memory deficits and result in disruptive consequences (McGarrity et al. 2017).

Posttranscriptional modifications of mRNA and microRNA molecules add a layer of complexity to gene expression, enabling context-specific regulation of translation, which is particularly crucial in neurons. Over two decades ago, it was hypothesized that cytoplasmic polyadenylation of synaptic mRNAs could be a major regulatory mechanism of local protein synthesis at synapses. The invertebrate cytoplasmic poly(A) polymerase TENT2 was shown to play a role in synaptic plasticity (Kwak et al., 2008). In mammals, its orthologue TENT2 (GLD2) has been proposed as a possible candidate for this function. However, there have been no reported brain-related physiological phenotypes of TENT2 knockout (KO) to date. Instead, the primary molecular phenotypes are associated with miRNA tailing rather than mRNA polyadenylation (Mansur et al., 2016; Mansur et al., 2021), despite its ubiquitous expression in the brain. Here we show that dysregulation of TENT2 RNA targets results in the imbalance in the inhibitory and excitatory neurotransmission manifested by increased excitability of *Tent2* KO neurons and has an effect on animal behavior. Our data thus establish a link between TENT2-dependent microRNA tailing and the balance of inhibitory and excitatory neurotransmission.

## Materials and Methods

### Generation of *Tent2* knockout mice

*Tent2* knockout mouse line in C57BL/6/Tar x CBA/Tar mixed background, were established using the CRISPR/Cas9 method in the Mouse Genome Engineering Facility (www.crisprmice.eu). The insertion of 21 bp cassette in exon 3 (insTAATAGTGAggatccCTAGAT after p.Leu86) intoduces 3xSTOP codons in the reading frame of TENT2 gene, as well as 1x STOP codons in +1 and +2 reading frames (Figure 1 A). Additionally, a unique BamHI restriction site for rapid and cost-effective genotyping was introduced in the insert. Based on the mouse genome (GRCm38/mm10 Assembly) a single guide RNA (sgRNA) was designed using an online CRISPR tool (http://crispr.mit.edu) and ordered from Synthego. The chosen sequence did not show any major off-targets but had high calculated efficiency.

Donor mice (B6CBAF1 Tar; C57BL/6/Tar x CBA/Tar) were injected first with 10 IU of PMSG (Pregnant Mare Serum Gonadotropin; Folligon, Intervet, Netherlands) and 48 hours later with 10 IU of hCG (Human Chorionic Gonadotropin; Chorulon, Intervet, Netherlands) to induce superovulation. Females were mated with males (B6CBAF1 Tar) immediately after hCG injection. Zygotes were collected from mated females 21–22 h post hCG injection. Zygotes, were microinjected into the cytoplasm using Eppendorf 5242 microinjector (Eppendorf-Netheler-Hinz GmbH) and Eppendorf Femtotips II capillaries with the following CRISPR cocktail: Cas9 mRNA IVT (25 ng/μl), sgRNA Tent2_KO IVT (12,5 ng/μl), oligo_Tent2_KO (1,33pM). After ∼2 hours after microinjection, culture microinjected zygotes were transferred into the oviducts of 0,5-day p.c. pseudo-pregnant females. Pups from the F0 generation were genotyped at around 4 weeks. The presence of insertion was confirmed by sequencing in the founder mouse and, after backcrossing, in N1 generation mice. All mice were bred and maintained in the animal house of the Faculty of Biology, University of Warsaw under a 12-h light/dark cycle with food and water available ad libitum. The animals were treated in accordance with the EU Directive 2010/63/EU for animal experiments.

## Genotyping

Pups were genotyped at 4 weeks of age. DNA from tail or ear tips was isolated with Genomic Mini kit (A&A Biotechnology). gDNA was amplified with Trap1_Seq1F/1R primer pair using Phusion HSII polymerase and HF buffer, then the amplicons were sequenced.

PCR cycling:

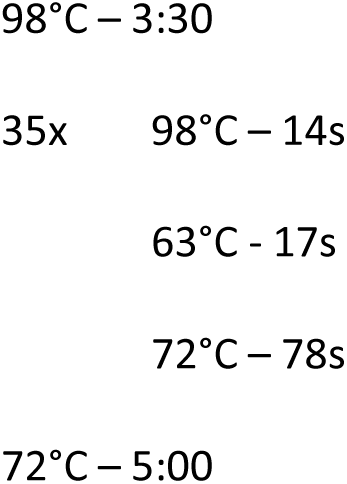

Primers used in the study:

sgRNA sequence: 5’ CGCTGCCGTTTACCATCAAG 3’

Tent2_Seq1F: 5’ ATGTGGTGACTCATTTTGGTAG 3’

Tent2_Seq1R: 5’ TAGAAAATTAGGTACTCCTGATCC 3’

oligo_Tent2_KO (ssDNA repair template): atttgcttgttttcagGAGAATAAGCGATGAAAAAGCTTTTCCTCTT<u>taatagtgaGGATCCCTagAt</u>gaGATGGTAAACG GCAGCGTTTCCATTCACCCCACCAAGAGCCAACTATAAT

### RNA isolation

Young adult male mice (6,5-week old) were sacrificed and the hippocampi were dissected. Total RNA was extracted using RNeasy Lipid Tissue Mini Kit (Qiagen). Next, 7 µg of total RNA from each sample was purified using KAPA Pure Beads (Roche) according to the manufacturer’s instructions and the quality and integrity of RNA was assessed on Agilent Bioanalyzer 2100 using Agilent RNA 6000 Pico Kit (Agilent, # 5067-1513). Purified RNA was further used for downstream analysis for total RNA sequencing and Nanopore direct RNA sequencing.

### Library preparation and total RNA-sequencing

RNA was ribodepleted and the libraries were prepared using KAPA RNA HyperPrep Kit with RiboErase (KAPA Biosystems, nr kat. 08098140702) according to the manufacturer’s recommendations. For library preparation 1µg of RNA was used, fragmented by 5 minutes incubation at 94°C. The library was enriched with 9 amplification cycles. The quality of the enriched library was verified using High Sensitivity D1000 Reagents (Agilent, nr kat. 5067-5585) on Agilent TapeStation 2200. The libraries’ concentration was estimated by qPCR means with KAPA Library Quantification Kit (Kapa Biosciences, Cat# KK4824), according to manufacturer’s instructions. These libraries were subsequently sequenced using an Illumina NovaSeq 6000 sequencing platform and NovaSeq 6000 SP Reagent Kit v 1.5 (200 cycles) (Illumina, Cat# 20040719) in 2×100nt pair-end mode with standard procedure according to manufacturer’s instructions with 1% control library Phix (.Illumina, Cat#. FC-110-3001).

### RNA Seq data analysis

The libraries were sequenced in pair-end mode on the NovaSeq 6000 Illumina platform. Obtained reads were adapter-clipped and quality-filtered with Atria (4.0.0)(Chuan et al. 2021)), then mapped to the reference mouse genome (GRCm39) with the STAR (Spliced Transcripts Alignment to a Reference) aligner (2.7.10a)( (Dobin et al. 2013)). Read counts were assigned to genes using featureCounts from the Subread (Liao, Smyth, and Shi 2014) package (2.0.6) with Gencode vM26 annotation file using default parameters. Multimappers and reads overlapping multiple features were excluded. Differential expression analysis was done with DESeq2 (1.34.0) (Love, Huber, and Anders 2014) with default settings. Fold change was corrected using apeglm algorithm (Zhu, Ibrahim, and Love 2019) Gene set enrichment analysis was performed in R (4.1.2) environment using g:Profiler (Raudvere et al. 2019) and GO-term analysis was done with ClusterProfiler (Wu et al. 2021). The results were visualised using ggplot2 package (Wickham, 2016)

### Nanopore direct RNA sequencing

Direct RNA-seq was performed as described by Bilska et al. (2020). Briefly, the 4.5-5 μg of total murine RNA was mixed with 150-200 ng of oligo(dT)25-enriched mRNA from Saccharomyces cerevisiae yeast, and spiked with 5 ng of *in vitro* transcribed poly(A) standards. Sequencing libraries were prepared with Direct RNA Sequencing Kit (Oxford Nanopore Technologies, SQK-RNA002) according to the manufacturer’s instructions. Sequencing was performed on a MinION device equipped with R9.4.1 RevD flow cells and controlled with MinKNOW software (Oxford Nanopore Technologies). Basecalling was performed with Guppy 6.0.0 (ONT).

### Poly(A) tail length profiling

Reads were mapped to Gencode vM26 (GRCm39) reference transcriptome with minimap2 2.17 (Li 2018) (-k 14 -ax map-ont --secondary = no) and processed with samtools 1.13 (Danecek et al. 2021) to exclude supplementary alignments and reads mapping to the reverse strand (samtools view -b -F 2320). The lengths of the poly(A) tails were estimated using the Nanopolish 0.13.2 polya function (Workman et al., 2019). In downstream analyses, only reads tagged by Nanopolish as “PASS” and “SUFFCLIP” were included. Statistical inference was performed using functions from the NanoTail R package (Krawczyk et al., 2024). The Wilcoxon signed-rank test was used to compare Poly(A) length distributions across different experimental conditions. Transcripts with a low number of supporting reads under each condition (<10) were excluded. P values were adjusted for multiple comparisons using the Benjamini-Hochberg method. Transcripts were considered as to having a significant change in poly(A) tail length if the length difference between the conditions was >=5nt and adjusted P value was less than 0.05. Gene set enrichment analysis was performed in R (4.1.2) environment using g:Profiler (Raudvere et al., 2019) and GO-term analysis was done with ClusterProfiler (Wu et al., 2021). The results were visualised using ggplot2 package (Wickham, 2016)

### miRNA isolation

Adult male mice (3-weeks old) were sacrificed and the hippocampi were dissected. miRNA was extracted using Direct-zol RNA MiniPrep kit (Zymo Research, cat. no.: R2050) according to the manufacturer instructions. The RNA quality was verified by Agilent Bioanalyzer 2100 with RNA 6000 Pico Kit (Agilent, nr kat. 5067-1513). miRNA-Seq libraries were prepared using TruSeqTM Small RNA Library Prep Kit (Illumina, nr kat. RS-200-0036). The size distribution of the final libraries was validated with Tape Station 2200 using High Sensitivity D1000 Reagents (Agilent, nr kat. 5067-5585). Libraries concentration was determined using qPCR with Kapa Library Quantification kit (Kapa Biosciences, nr kat. KK4824). For all above procedures manufacturer’s protocols were used.

### miRNA sequencing

The sequencing was performed using Illumina NovaSeq 6000 with the NovaSeq 6000 SP Reagent Kit v 1.5 (200 cycles) (Illumina, nr kat. 20040719), generating 2 × 100 pair-end reads using the manufacturer’s standard protocols with 1% addition of the control library Phix (Illumina, nr kat. FC-110-3001).

Raw sequencing reads were trimmed to remove excessive adapter and sequencing primer sequences using the BBDuk, a part of BBTools (BBMap – Bushnell B. – sourceforge.net/projects/bbmap/). Paired reads for each sample were processed together and specific parameters were applied (k=23 mink=11 hdist=1 ktrim=r minlen=17 tpe). For miRNA expression levels only R1 reads were used. The miRNA expression levels and composition of the 3’ end of isomiRs were analyzed using QuagmiR (Bofill-De Ros et al., 2019). The QuagmiR was applied with searching for 5’ end variation filtering turned off and with specific custom parameters (min_ratio: .1 min_read: 9 edit_distance_3p: 3). Mature miRNA sequences, primiRNA sequences and annotations of Mus musculus were from miRBase v22. The miRNA expressions and isomiR composition can be found in Supplementary Figure S1.

In the downstream analysis we classified isomiRs as: (i) canonical (fully complementary to annotated mature miRNA sequence), (ii) trimmed (shorter than the annotated mature miRNA sequence), (ii) non templated (NT) tail (longer than the annotated mature miRNA sequence and tail sequence does not match genomic sequence), (iv) ambiguous tail (longer than the annotated mature miRNA sequence and tail sequence match genomic sequence, or shorter than the annotated mature miRNA sequence and tail sequence does not match genomic sequence). NT-tail was also analyzed in terms of tail length and specific sequences focusing on tails containing mono or combinations of adenosines and uridines.

All statistical analysis of isomiR classes and NT tail were performed with R custom scripts. The Wilcoxon test was used to determine *P*-values, and values < 0.05 were considered statistically significant. The differential expression of isomiRs was performed with use of the DESeq2 R package (Love et al., 2014) with default settings. As cutoff for significant differentially expressed isomiRs a threshold of adj. *P*-value <0.05 and |log2(fold change)| > 1 was set. All visualizations were performed with the ggplot2 R package [Wickham H (2016). *ggplot2: Elegant Graphics for Data Analysis*. Springer-Verlag New York. ISBN 978-3-319-24277-4, https://ggplot2.tidyverse.org.]

### Patch clamp electrophysiology in acute brain tissue slices

3-5 weeks old mice were anesthetized with Isoflurane and decapitated. Brains were isolated and transferred into ice-cold cutting artificial cerebrospinal fluid (ACSF) containing 87 mM NaCl, 2.5 mM KCl, 1.25 mM NaH2PO4, 25 mM NaHCO3, 0.5 mM CaCl2, 7 mM MgSO4, 20 mM D-glucose, 75 mM sacharose equilibrated with carbogen (5% CO2/95% O2). The brain was cut into 2 hemispheres, and 350 μm-thick horizontal brain slices were cut in ice-cold cutting ACSF on a 5100mz vibratome (Campden Instruments, Loughborough, England). Slices were then incubated for 15 minutes in cutting ACSF at 32°C. Next, the slices were transferred to recording ACSF containing: 125 mM NaCl, 2.5 mM KCl, 1.25 mM NaH2PO4, 25 mM NaHCO3, 2.5 mM CaCl2, 1.5 mM MgSO4, 20 mM D-glucose equilibrated with carbogen and incubated for minimum 1 hour at RT. Patch-clamp recordings were recorded in a submerged chamber perfused with recording ACSF at 3 mL/min in RT. Patch pipettes (resistance 4 to 6 MΩ) were pulled from borosilicate glass (WPI, 1B120F-4) with a micropipette puller (Sutter Instruments, P-1000) and filled with internal solution containing 95 mM potassium gluconate, 6 mM KCl, 2 mM NaCl, 0.5 mM EGTA, 20 mM HEPES, 4 mM MgATP, 0.4 mM NaGTP, and 10 mM Na2 phosphocreatine (pH 7.4). For each slice, the measurements were done in stratum pyramidale of CA1. Firing pattern was recorded in the current-clamp mode by injection of depolarizing current stimulus. Recordings were acquired and digitized with Dual IPA amplifier (Sutter Instruments, Novato, CA, USA) and analyzed with Clampfit 10.7 software (Molecular Devices, San Jose, CA, USA) and AxoGraph 1.7.4 software (developed by Dr John Clements).

### Patch-clamp electrophysiology in primary hippocampal neuronal cultures

Whole-cell patch-clamp recordings in voltage-clamp mode were performed in 21-22 DIV in hippocampal cultures using borosilicate patch pipettes (4–6 MΩ) filled with one of the intracellular solutions described below. The external solution contained 120 mM NaCl, 5 mM KCl, 2 mM CaCl2, 1 mM MgCl2, 20 mM glucose, and 10 mM HEPES (pH 7.4, adjusted with NaOH). The mEPSCs were recorded at a holding potential of − 60 mV after bath application of 10 μM gabazine and 1 μM tetrodotoxin (TTX). The internal solution contained 95 mM potassium gluconate, 6 mM KCl, 2 mM NaCl, 0.5 mM EGTA, 20 mM HEPES, 4 mM MgATP, 0.4 mM NaGTP, and 10 mM Na2 phosphocreatine (pH 7.4). At the end of recording, 20 μM AMPA/kainate receptor antagonist 6,7-dinitroquinoxaline-2,3-dione (DNQX) was used to confirm the origin of the recorded mEPSCs. The mIPSCs were recorded at a holding potential of − 60 mV in the presence of 1 μm TTX and 20 μM DNQX with an internal solution containing 140 mM CsCl, 2mM NaCl, 10mM HEPES, 5 mM EGTA, 2mM MgCl2, 2mM Na2-ATP (pH 7.4).

After the end of recording, 10 μM GABAA receptor antagonist gabazine was used to confirm the origin of the recorded mIPSCs. Extracellular solution was perfused at 3 mL/min. All recorded signals were low-pass filtered at 5 kHz using the filter built into the Dual IPA patch-clamp amplifier, digitized at 20 kHz (Dual IPA amplifier, Sutter Instruments, Novato, CA, USA), and acquired with the SutterPatch software (Sutter Instruments, Novato, CA, USA. The series resistance (Rs) was estimated from the response to a hyperpolarizing voltage step (−5 mV). The recordings in which series resistance was > 20 MΩ were rejected. All electrophysiological data were analyzed with Clampfit 10.7 software (Molecular Devices, San Jose, CA, USA) and AxoGraph 1.7.4 software (developed by Dr John Clements).

### Behavioral assays

Adult male WT and *Tent2* KO mice were used. Experiments were performed sequentially in the following order: marble burying, elevated plus maze, open field test. A cohort of: WT, n=9 and KO, n=8 mice was used. The contextual fear conditioning was performed with another three separate cohorts of animals (WT n=4, n=7, n=11 and KO n=4, n=6, n=8). Mice were habituated to the room and the experimenter, assays were conducted during the light phase of the light/dark cycle. All the apparatus used were thoroughly cleaned with 70% ethanol between animals. Mouse behavior was video recorded and analyzed using EthoVision XT (Noldus).

### Marble burying

The test was conducted in polypropylene box (43 cm length, 26 cm width, 31 cm height) filled with 5cm depth of fresh bedding. Animals were habituated to the box two days for ∼1 h each day. On the third day fifteen clear glass marbles (2 cm diameter) were placed evenly spaced in three columns on the surface of the bedding. Animals were placed separately onto the central area of the box and allowed to freely explore the new environment. After 30 min of testing, mice were placed back to their home-cage and number of marbles that were buried to 2/3rd of their depth were counted manually. Whole experiments were videotaped by the overhead mounted camera and analyzed with the EthoVision XT tracking system (Noldus, Wageningen, NL). Each mouse was given fresh bedding and marbles were previously cleaned with 70% ethanol and water to eliminate any olfactory cues. All experiments were performed in a darkened lighting conditions (4-5 lux).

### Elevated plus maze

Testing was performed in a metal plus-maze apparatus elevated to a height of 50 cm (AnimaViVari). The apparatus consisted of two open arms and two closed arms both of the same dimensions (35 x 5 cm), with gray PCV walls 15 cm high and arranged so that both the open and closed arms faced each other. The testing was done in a dimly lit (30 Lx at the maze level) area surrounded with non-transparent, grey curtains in order to limit any additional spatial stimuli. Mice were placed at the intersection of the open and closed arms, and mouse behavior was recorded by camera during a 5-min test interval and analyzed with the EthoVision XT tracking system (Noldus, Wageningen, NL). The following parameters were calculated - in open and closed arms: number of entries, total time spent, distance moved and movement duration; in the central platform: time spent; in the whole arena: total distance moved, total movement duration.

### Open field test

Testing was performed in a plexiglass box (dimensions 64 x 64, height 32 cm). Mice were placed in the middle of the arena and animal’s movements were recorded for 15 minutes by a camera set up above the arena. The video was analyzed using a EthoVision XT video-tracking system (Noldus, Wageningen, NL) to extract behavioral data. Two zones, the center zone (inner 50 cm diameter circle) and the border zone (remaining 6 cm border), were defined according to previously obtained criteria. The following parameters were taken into analysis - time spent, distance moved and movement duration. The arena was cleaned after each animal with a 70% solution of ethanol and dried.

### Contextual Fear Conditioning

Mice were trained in a Med Associates Inc. Fear Conditioning Chamber (St Albans, USA) connected to a computer running Video Freeze software. Thirty minutes before training mice were brought to the room with conditioning chamber to acclimatize. Chambers were cleaned with 70% ethanol and the paper towels soaked with ethanol were placed under the metal grid. Mice were placed in the chamber on a metal grid platform and after 148 seconds of habituation, received three electric shocks (US, 2 s, 0.7 mA) with 90 s intervals -the training lasted 6 min. The animals were taken out from the experimental chamber 30 s after the last shock and put in the homecage. Between animals the cage was cleaned with 70% ethanol solution and a fresh paper towel was placed. The fear memory of the context, defined as a level of freezing in the context, was assessed for 5 min in the same experimental chamber 1h, 24 h and 48 h after training. All animals were trained, tested and sacrificed during the light phase of animals’ day (between 09.00 and 16.00 h). The training and testing times were counterbalanced between the groups. Animals were sacrificed after the last test session, hippocampi were dissected and frozen on dry ice.

### Quantitative determination of amino acid profile

Frozen hippocampi were lyophilized using Alpha 2-4 LD Plus lyophilizer (Martin Christ). Amino acid levels were measured using High-performance liquid chromatography coupled with tandem mass spectrometry (LC-MS/MS) in a commercial laboratory (Masdiag, Warsaw, Poland).

## Data Availability

The raw sequencing data (fast5 files) were deposited at the European Nucleotide Archive (ENA) (https://www.ebi.ac.uk/ena/browser/home) under the project numbers (PRJEB76814 and PRJEB75356) and accession numbers and sample numbers listed in Supplementary Table 1. The sRNA-Seq data for this study have been deposited in the European Nucleotide Archive (ENA) at EMBL-EBI under accession number PRJEB76999 (https://www.ebi.ac.uk/ena/browser/view/PRJEB76999).

## Acknowledgements

The research leading to these results was funded by the Norwegian Financial Mechanism 2014 -2021, no. UMO-2019/34/H/NZ3/00733. IIMCB core facilities (IN-MOL-CELL infrastructure) were funded by the European Union and co-financed under the European Funds for Smart Economy 2021-2027 (FENG).

